# Conjugation of palbociclib with MHI-148 has an increased cytotoxic effect for breast cancer cells and an altered mechanism of action

**DOI:** 10.1101/2021.11.02.466693

**Authors:** Euphemia Leung, Petr Tomek, Moana Tercel, Jóhannes Reynisson, Thomas Park, Elizabeth Cooper, William Denny, Peter Choi, Jiney Jose

## Abstract

The CDK4/6 inhibitor palbociclib, combined with endocrine therapy, has been shown to be effective in postmenopausal women with estrogen receptor positive, HER2-negative advanced or metastatic breast cancer. However, palbociclib is not as effective in the highly aggressive triple-negative breast cancer that lacks sensitivity to chemotherapy or endocrine therapy. We hypothesized that conjugation of the near-infrared dye MHI-148 with palbociclib can produce a potential theranostic in triple-negative as well as estrogen receptor positive breast cancer cells. In our study, the conjugate was found to have enhanced activity in all mammalian cell lines tested *in vitro*. However, the conjugate was cytotoxic and did not induce G1 cell cycle arrest in breast cancer cells suggesting the mechanism of action differed from the parent compound palbociclib. The study highlights the importance of investigating the mechanism of conjugates of near-infrared dyes to therapeutic compounds as conjugation can potentially result in a change of mechanism or target, with an enhanced cytotoxic effect in this case.

## Introduction

Breast cancer is a worldwide health concern with over two million new cases in 2020 (https://www.who.int/news-room/fact-sheets/detail/breast-cancer). Significant advances have been made in understanding different breast cancers and several molecular subtypes of breast cancer have been characterized.^1^ This knowledge has accelerated the development of new therapies to target molecular alterations that drive tumor cell growth. Abnormal proliferation with dysregulation of normal cell cycle control are common to all cancer types.^2^ Cell cycle progression plays a crucial role in cell proliferation, and alterations in cell cycle regulators have been acknowledged as a hallmark of cancer.^2^ Cyclin-dependent kinases, CDK4 and CDK6, drive cell cycle progression from G0 or G1 phase into S phase where DNA replication occurs.^3^ Therefore, CDK4/6 are appealing targets for novel cancer therapeutics.^4^

Palbociclib (a selective CDK4/6 inhibitor) has been approved by the Food and Drug Administration for postmenopausal women with HR (hormone-receptor)-positive, HER2 (human epidermal growth factor receptor 2)-negative advanced or metastatic breast cancer in combination with an aromatase inhibitor or fulvestrant, and recently the approval has been expanded to include male breast cancer.^5^ Although palbociclib has high selectivity, several relevant adverse effects including neutropenia have been reported.^6^ To increase efficacy of palbociclib and reduce adverse effects, one strategy is to conjugate the drug to an antibody or small molecule that can enhance intracellular drug concentrations in cancer cells relative to non-cancerous cells, increasing therapeutic indices.^7^

Theranostics is a combination of the term therapeutics and diagnostics.^8^ Theranostics are small molecules or nano-size compounds serving for both therapy and diagnosis. They have attracted much attention as they can play a vital role in reducing the side effects and evaluating the therapeutic efficiency of a prodrug *in vivo*.^9^ Heptamethine cyanine dyes (HMCDs) are a class of near-infrared fluorescent (NIRF) compounds that have recently emerged as promising agents for drug delivery with tumor-targeting properties.^10^ Initially explored for their use in imaging neoplasms, their tumor-targeting properties and their tumor selectivity is primarily attributed to their uptake by isoforms of organic anion transporting polypeptides (OATPs) which are overexpressed in cancer tissues.^11,12^ A particular heptamethine cyanine dye (MHI-148) was reported to readily form an adduct with albumin,^13^ and MHI-148 can be imported via albumin receptors which are over-expressed on cancer cells.^14^ MHI-148 has previously been conjugated to kinase inhibitors (e.g. dasatinib^15^) with the goal of selectively delivering them to solid tumors.^16,17^ Since MHI-148 was reported to have tumor-targeting capability,^18^ it was used as the NIRF agent in our study. Therefore, we resynthesized the MHI-148 palbociclib conjugate, hereafter referred to as MHI-palbociclib, as reported in a patent (US 2019/0343958) to examine its effect and explored the mechanism of action of the conjugate in oestrogen receptor positive MCF-7 and triple-negative MDA-MB-231 human breast cancer cell lines.

## Materials and Methods

### Synthesis of MHI-palbociclib

MHI-palbociclib was designed to connect palbociclib to MHI-148 via the solvent-exposed piperazine group of the kinase inhibitor, according to a previously published protocol (https://www.freepatentsonline.com/20190343958.pdf)

### Absorbance and fluorescence spectra measurements

The absorbance and fluorescence emission spectra of up to 7 three-fold serial dilutions of MHI-148 and MHI-palbociclib in DMSO ranging from 10^2^ to 10^−2^ μM were acquired at 25°C. We used DMSO instead of a more physiologically relevant aqueous buffer for dissolving the compounds because MHI-palbociclib rapidly precipitates in aqueous environments in high concentration.

a. Absorbance spectra of the test compound solutions (200 μL) plated in a 96-well polystyrene flat-bottom transparent microplate were acquired on a quad monochromator-equipped microplate reader Enspire 2300 (Perkin-Elmer; Singapore) from 650 nm to 900 nm in 1 nm increments.
b. Fluorescence emission spectra of the compound solutions in a 2 mL semi-micro quartz cuvette were measured on a FP-8600 spectrofluorometer (JASCO; Japan) between 790 nm and 850 nm in 1 nm increments at the excitation wavelength of 780 nm. The fluorescence emission was measured at high sensitivity of the instrument, excitation and emission bandwidth of 5 nm, 50 ms response time and scan speed of 1000 nm per minute.

### Cell lines

All cell lines were passaged in αMEM supplemented with 5% fetal bovine serum (FBS), and insulin/transferrin/selenium supplement (Roche) without antibiotics for less than 3 months from frozen stocks confirmed to be mycoplasma-free. The human breast cancer oestrogen receptor positive MCF-7 and triple-negative MDA-MB-231 cell lines, and the human embryonic kidney cell line HEK293 were purchased from the American Type Culture Collection (ATCC). The Chinese hamster ovary CHO cell lines, and their DNA repair genotypes and origins, were as follows: 51D1^19^ (rad51d knockout) and 51D1.3^19^ (51D1 complemented with CHO Rad51d). All cell lines tested negative for mycoplasma contamination (PlasmoTestTM - Mycoplasma Detection kit, InvivoGen, San Diego, CA).

### Cell proliferation assays and viability assay

As described previously,^20^ 3000 cells per well were seeded in 96-well plates and incubated in the presence of varying concentrations of drugs for three days. Cell proliferation was measured by thymidine uptake assay, cell growth by sulforhodamine B (SRB) colorimetric assay^21^ and cell viability by WST-1 assay as previously described. ^22^

Briefly, for the proliferation assay,^23^ [^3^H]-thymidine (0.04 μCi) was added to each well and plates were incubated for 5 h, after which the cells were harvested onto glass–fiber filters using an automated TomTec harvester. Filters were incubated with Betaplate Scint and [^3^H]-thymidine incorporation was measured in a Trilux/Betaplate counter. Cell proliferation was determined by the percentage incorporation of [^3^H]-thymidine.

The SRB colorimetric assay, which is based on the measurement of cellular protein content, was used to measure cell density.^21^ After drug treatment for 3 days, cells were fixed with 10% (wt/vol) trichloroacetic acid and stained for 30 min, and the excess dye was removed by washing repeatedly with 1% (vol/vol) acetic acid. The protein-bound dye was dissolved in Tris base solution (10 mM) for optical density determination at 510 nm using a microplate reader. Optimal cell densities were previously determined to select initial cell densities that ensured that cells were in logarithmic phase for the experiments.

For viability, superoxide dismutase activity was measured using a water-soluble tetrazolium salt (WST-1) (purchased from Roche) after drug treatment for 3 days. The stable tetrazolium salt WST-1 is cleaved to a soluble formazan by a complex cellular mechanism that occurs primarily at the cell surface. This bioreduction is largely dependent on the glycolytic production of NAD(P)H in viable cells. Therefore, the amount of formazan dye formed directly correlates with the number of metabolically active cells in the culture.

All experiments were carried out using duplicate wells in three independent experiments.

### Measurement of DNA content for cell cycle analysis

As previously described,^24^ cells (1 × 10^6^ cells) were grown in six-well plates and incubated with drugs for 24 h. Cells were harvested, washed with 1% FBS/phosphate-buffered saline (PBS), resuspended in 200 μl of PBS, fixed in 2 ml of ice-cold 100% ethanol, and stored overnight at −20°C. The cells were washed and resuspended in 1 ml of 1% FBS/PBS containing RNase (1 μg/ml) and propidium iodide (PI) (10 μg/ml) for 30 min at room temperature. DNA content was determined using forward scatter (FSC) intensity by PI staining based on a total of 25,000 acquired events on a BD Biosciences Accuri C6 flow cytometer and cell cycle distribution was analysed using FlowJo version 10.7.2.

### Propidium iodide staining

Cells (1 × 10^6^) were incubated with drugs for 24 h. Cells were next incubated with 50 μg/ml PI in growth media for 15 min at room temperature and protected from light. Cell death was monitored by fluorescence staining of DNA by PI and analysed by fluorescent microscopy using Floid imaging station (460× magnification).

### Confocal imaging

MCF-7 cells were treated with 20 μM MHI-palbociclib at 37°C for 2 h and MitoTracker green for 30 min before imaging. Images were captured using a Zeiss LSM 800 Airyscan confocal microscope. Single optical sections of the cells were imaged with a 10×/0.45 NA Plan Apochromat objective lens. Red fluorescence was acquired using a 561nm diode-pumped laser with an emission range of 570-700 nm. Green fluorescence was captured using a 488 nm diode laser with an emission range of 493-585 nm. The field of view was 638.90 × 638.90 microns.

### Chemical space

The Scigress version FJ 2.6 program^25^ was used to build the compound structures and derive their molecular descriptors; the MM3^26-28^ force field was applied to identify the global minimum using the CONFLEX method^29^ followed by structural optimization. The Log P values were derived according to Ghose et al.^30^ and the solvent accessible surface (SAS) area was calculated at an optimized geometry in water; the water geometry is from optimization using MO-G with PM6 parameters and the Conductor like Screening Model (COSMO).^31^

### Statistical analysis

Results are presented as mean ± SEM. Unpaired t-tests were used for comparison between two groups.

## Results

### Physicochemical properties of the NIRF dye MHI-148 and its conjugate with palbociclib

Palbociclib^32^ was conjugated to MHI-148^33^ via the solvent-exposed piperazine group (Figure 1). The conjugation of palbociclib to MHI-148 did not significantly change the absorbance or fluorescence properties of the parent MHI-148 dye in DMSO solution (Figure 1).

**Figure 1.**
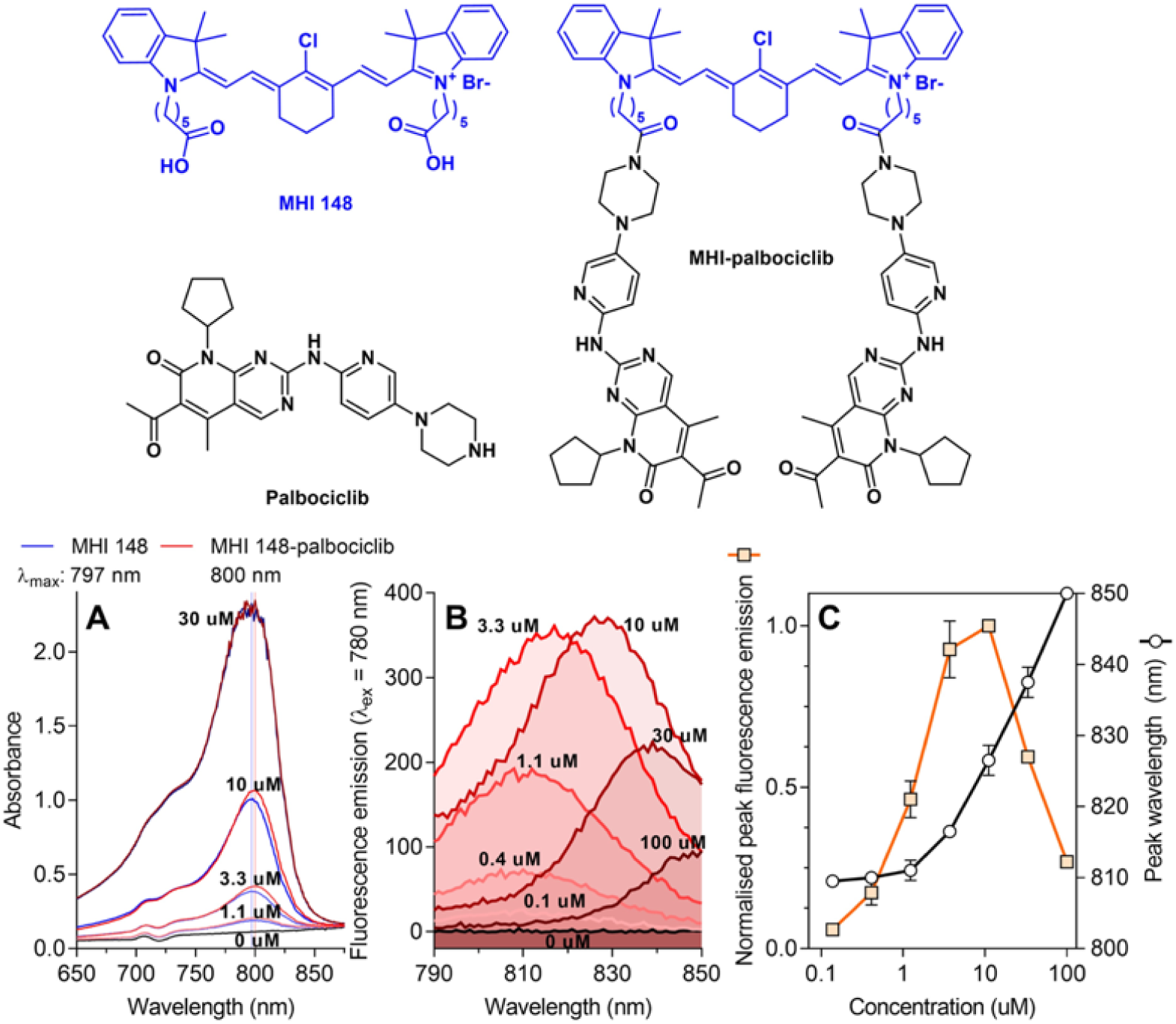
The conjugation of palbociclib to MHI-148 does not significantly influence the absorbance and fluorescence properties of the parent MHI-148 dye. **(A)** Absorbance spectra of MHI-148 and MHI-palbociclib in DMSO. A representative experiment of two independent repeats. Vertical lines denote λ_max_ which indicates the wavelength at the peak maximum. **(B)** Fluorescence spectra of MHI-palbociclib in DMSO. Each fluorescence emission spectrum represents an average of three measurements. Only MHI-palbociclib is displayed as MHI-148 showed essentially identical normalised spectra. **(C)** Impact of fluorophore concentration on fluorescence intensity and peak wavelength. Each data point represents a mean and standard deviation of respective values for MHI-148 and MHI-palbociclib obtained from data in panel B. Increasing MHI-palbociclib’s concentration red-shifts the fluorescence and induces self-quenching.

### Effect on cell proliferation, growth and viability

The effect of MHI-palbociclib was compared to palbociclib and MHI-148 in breast cancer and non-cancerous cell lines where cell proliferation (measured by [^3^H]-thymidine incorporation), cell growth (measured by SRB assay), and cell viability (measured by WST-1 assay) were examined (Figure 2).

**Figure 2.**
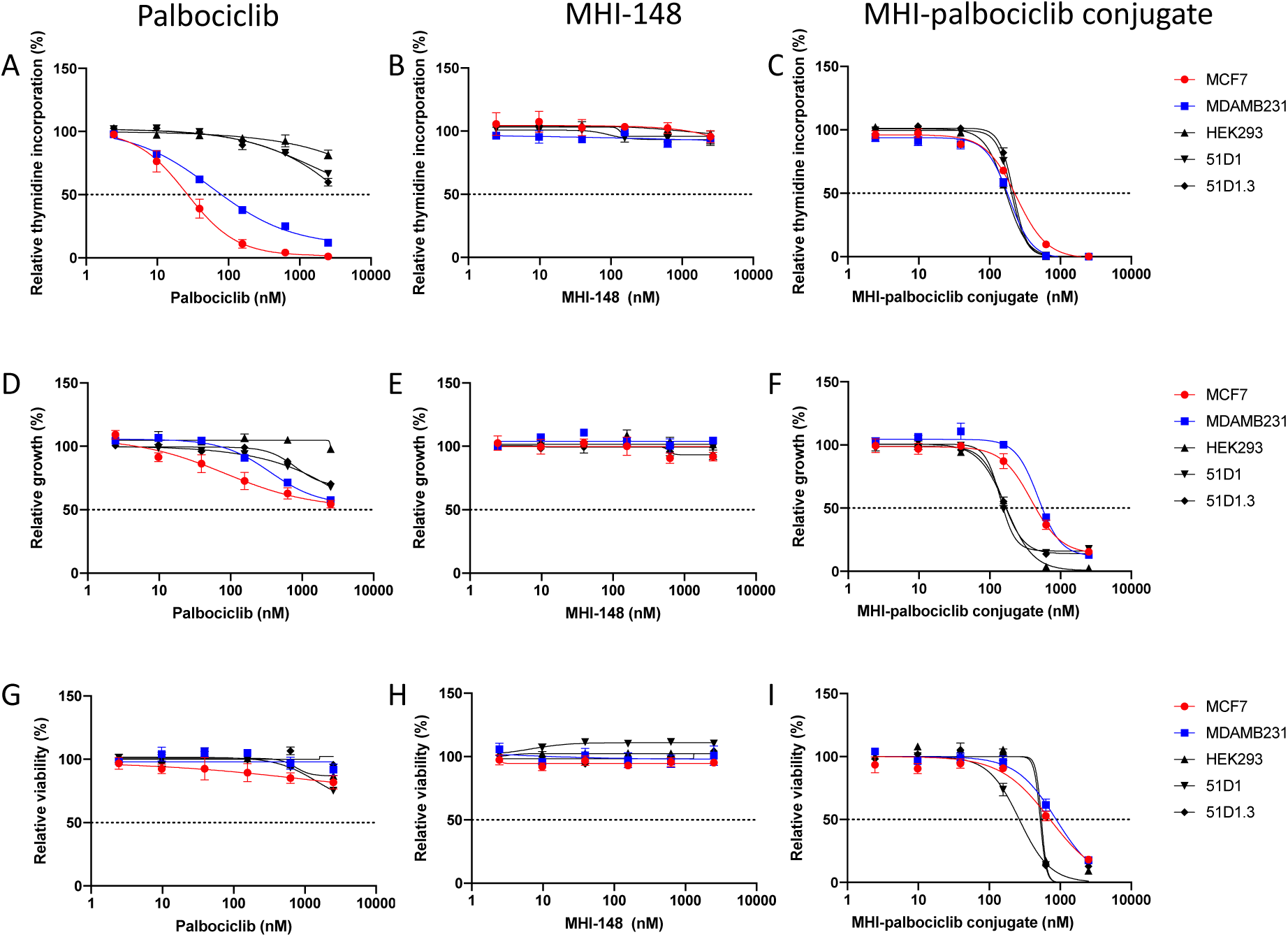
Effect of palbociclib, MHI-148 and MHI-palbociclib on **(A-C)** cell proliferation ([^3^H]-thymidine incorporation assay), **(D-F)** cell growth (SRB assay) and **(G-I)** cell viability (WST-1 assay). Effects on breast cancer cell lines and non-cancerous cell lines were measured after 3 days treatment with the compounds. Data presented is an average of at least three independent experiments.

The breast cancer oestrogen receptor-positive MCF-7 cells showed higher sensitivity than the triple-negative MDA-MB-231 cells towards palbociclib in the proliferation assay (Figure 2A), while the non-cancerous cells showed significantly increased resistance (significantly higher IC_50_ values) as compared to the breast cancer cell lines (Table 1 and Supplementary Table S1). Our data is similar to the previously reported palbociclib IC_50_ for MCF-7 and MDA-MB-231 (148 ± 25.7 and 432 ± 16.1 nM, respectively) determined using a different proliferation assay (a 2.5-fold versus 2.9-fold difference comparing the IC_50_ of the two cell lines using our data and the data from Finn et al., respectively).^34^ As a highly selective CDK4/6 inhibitor, palbociclib is expected to have cytostatic effects by causing cell cycle arrest at the G1/S checkpoint which leads to disruption of cancer cell proliferation. ^35,36^ We therefore confirmed that the viability (measured by WST-1 assay) of the tested cell lines was not affected by palbociclib treatment (Figure 2G). Increased inhibition (measured by SRB assay) was observed in all the cell lines tested (Figure 2D).

**Table 1.**
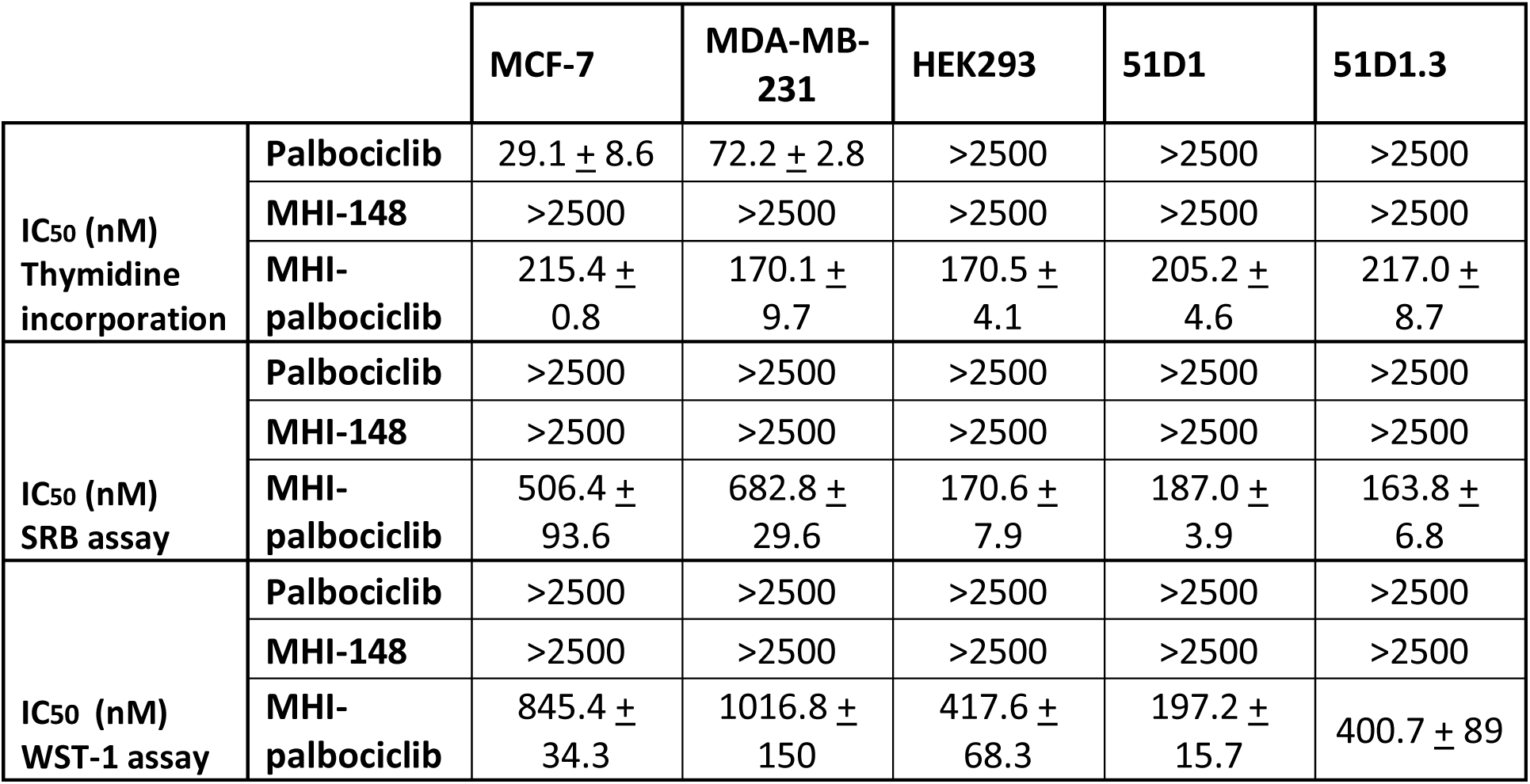
IC_50_ values of palbociclib, MHI-148 and MHI-palbociclib in breast cancer MCF-7 and MDA-MB-231 cell lines, and non-cancerous HEK293, 51D1 and 51D1.3 cell lines.

|  |  | MCF-7 | MDA-MB-231 | HEK293 | 51D1 | 51D1.3 |
| --- | --- | --- | --- | --- | --- | --- |
| IC<sub>50</sub> (nM)<br>Thymidine<br>incorporation | Palbociclib | 29.1 ± 8.6 | 72.2 ± 2.8 | >2500 | >2500 | >2500 |
|  | MHI-148 | >2500 | >2500 | >2500 | >2500 | >2500 |
|  | MHI-palbociclib | 215.4 ± 0.8 | 170.1 ± 9.7 | 170.5 ± 4.1 | 205.2 ± 4.6 | 217.0 ± 8.7 |
| IC<sub>50</sub> (nM)<br>SRB assay | Palbociclib | >2500 | >2500 | >2500 | >2500 | >2500 |
|  | MHI-148 | >2500 | >2500 | >2500 | >2500 | >2500 |
|  | MHI-palbociclib | 506.4 ± 93.6 | 682.8 ± 29.6 | 170.6 ± 7.9 | 187.0 ± 3.9 | 163.8 ± 6.8 |
| IC<sub>50</sub> (nM)<br>WST-1 assay | Palbociclib | >2500 | >2500 | >2500 | >2500 | >2500 |
|  | MHI-148 | >2500 | >2500 | >2500 | >2500 | >2500 |
|  | MHI-palbociclib | 845.4 ± 34.3 | 1016.8 ± 150 | 417.6 ± 68.3 | 197.2 ± 15.7 | 400.7 ± 89 |

MHI-148 did not show any inhibitory effect on proliferation, growth, or viability in any of the cell lines tested (Figure 2B, E and H).

In contrast, MHI-palbociclib displayed strong inhibitory effects on proliferation, growth and viability (Figure 2C, 2F and 2I, Table 1). Unlike the increased sensitivity to palbociclib in breast cancer cells, both breast cancer cells and non-cancerous cells (HEK293, CHO variants 51D1 and 51D1.3) showed similar sensitivity to MHI-palbociclib in cell growth (Figure 3C). Surprisingly, MHI-palbociclib significantly decreased cell viability as compared to palbociclib (Figure 2I and Table 1).

**Figure 3.**
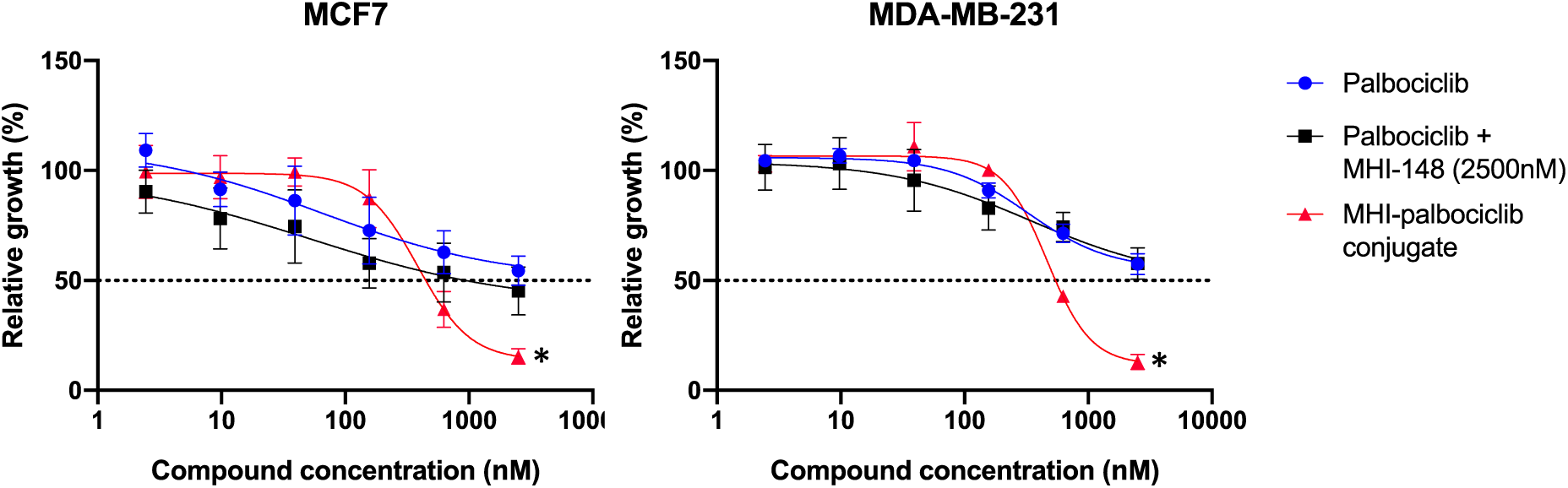
Inhibition of relative growth by MHI-palbociclib is significantly higher than the parent compound palbociclib, or the combination of palbociclib with unconjugated MHI-148, in breast cancer MCF-7 and MDA-MB-231 cell lines. Effects on breast cancer cell lines were measured after 3 days exposure to the compounds. Data presented is an average of at least three independent experiments.

The growth inhibitory effect of MHI-palbociclib at 2.5 μM was significantly greater than that of the combination of individual unconjugated compounds (palbociclib and MHI-148) in MCF-7 (T-test, *p* = 0.002) and MDA-MB-231 (T-test, *p* = 0.003) (Figure 3), indicating the enhanced effect was not due to co-incubation of the unconjugated NIRF dye MHI-148 and palbociclib.

### Cell cycle arrest

The inhibitory effect of the cytostatic compound palbociclib on proliferation (measured by thymidine uptake) were many fold higher than growth (measured by SRB) and viability (measured by WST-1) (Figure 2). The main basis for inhibition of proliferation was the induction of cell cycle arrest by palbociclib.^3^ Whereas MHI-palbociclib exhibited similar IC_50_ on cell proliferation, growth and viability, suggesting the characteristic is more similar to a cytotoxic compound. Since palbociclib treatment is known to induce G1 cell cycle arrest in MDA-MB-231 cells,^35^ we assessed whether MHI-palbociclib shares the same function. Recapitulating previously reported evidence, palbociclib treatment resulted in a prominent G1 cell cycle arrest in MDA-MB-231 cells versus untreated cells (87% vs 48%, respectively). A significant reduction in G1 cell cycle arrest in MHI-palbociclib treated cells (55%) as compared to palbociclib treated cells (87%) was observed (T-test, *p* < 0.0001) (Figure 4), further suggesting that the mode of action of MHI-palbociclib in this cell line is different to that of palbociclib.

**Figure 4.**
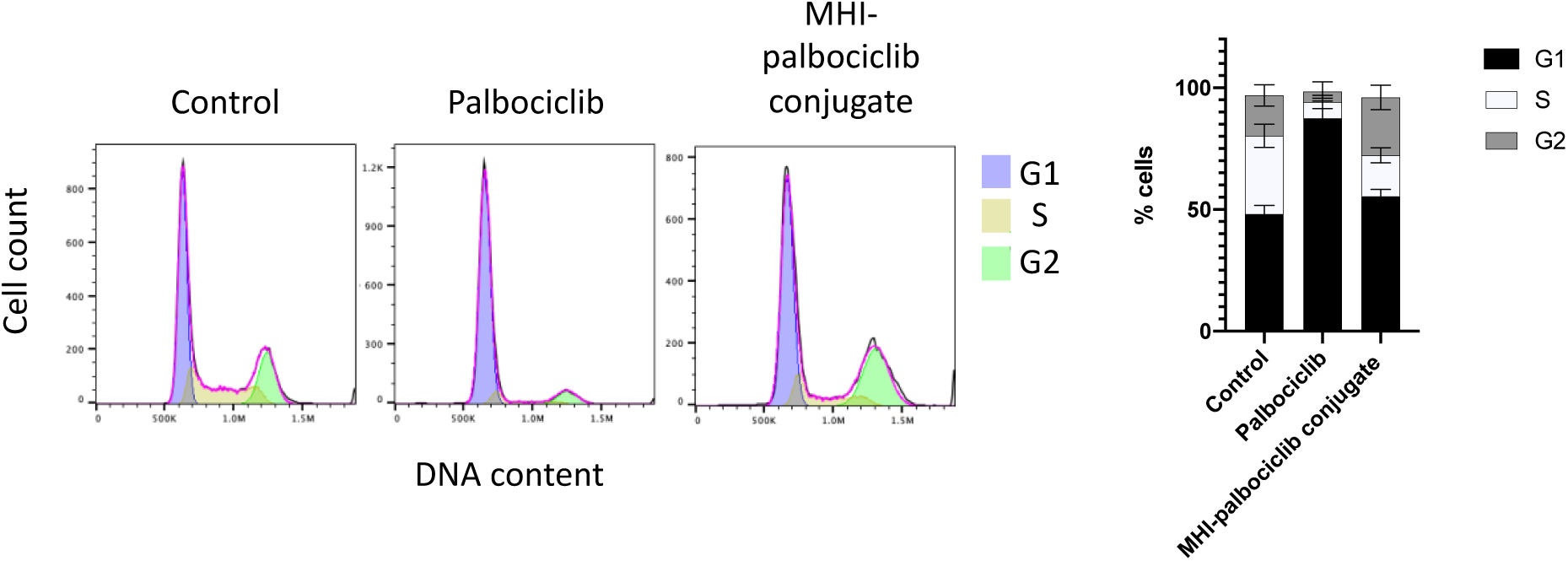
The cell cycle proportions of MDA-MB-231 breast cancer cells after exposure to 500 nM palbociclib or MHI-palbociclib for 24 h, followed by flow cytometry cell cycle analysis. Control: no drugs administered. Representative cell cycle profiles are shown on the left, and the average result of three independent experiments are shown in the bar graph.

### Cytotoxic property of MHI-palbociclib

To ascertain typical features of dying cells, the membrane-impermeable dye propidium iodide (PI) was used to show binding to DNA upon membrane damage, which occurs at late apoptotic or early necroptotic events.^37^ We demonstrated that increased PI staining and reduced cell number were observed in MHI-palbociclib-treated cells but not palbociclib-treated or untreated cells, supporting the cytotoxic effect of MHI-palbociclib (Figure 5).

**Figure 5.**
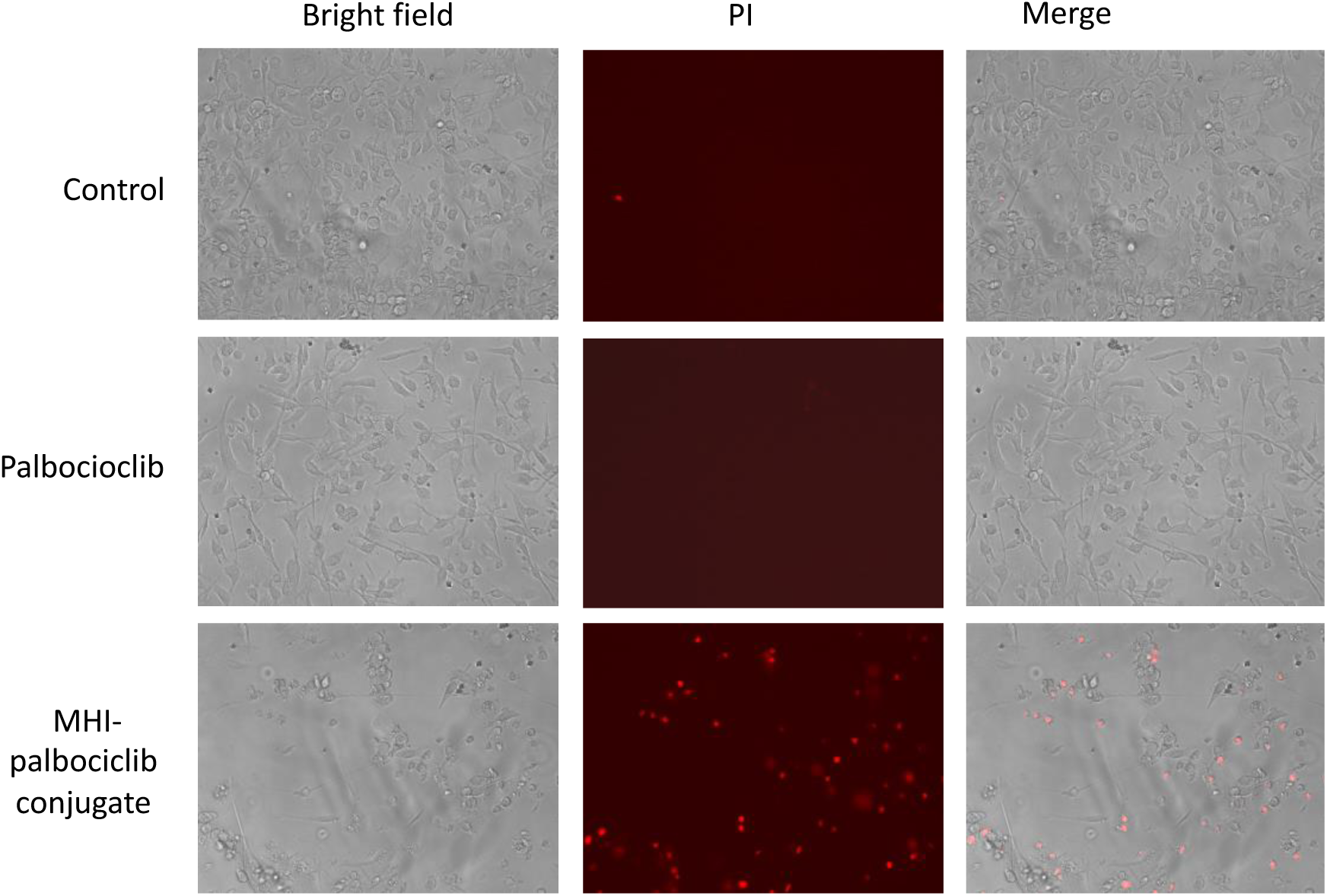
Cytotoxicity in MDA-MB-231 cells after MHI-palbociclib treatment is significantly greater than after treatment with palbociclib. After exposure to 1 μM of indicated compound for 24 h, dying cells were stained by propidium iodide (red). Photographs were taken by Floid imaging station (460× magnification). Representative images of three independent experiments are shown.

### Mitochondrial Localization

We investigated the subcellular localization of MHI-palbociclib in the MCF-7 breast cancer cell line. Merged images revealed an overlap of the cytoplasmic staining of MHI-palbociclib with a fluorescent marker of mitochondria (Figure 6), indicating the targeting potential of MHI-palbociclib to mitochondria.

**Figure 6.**
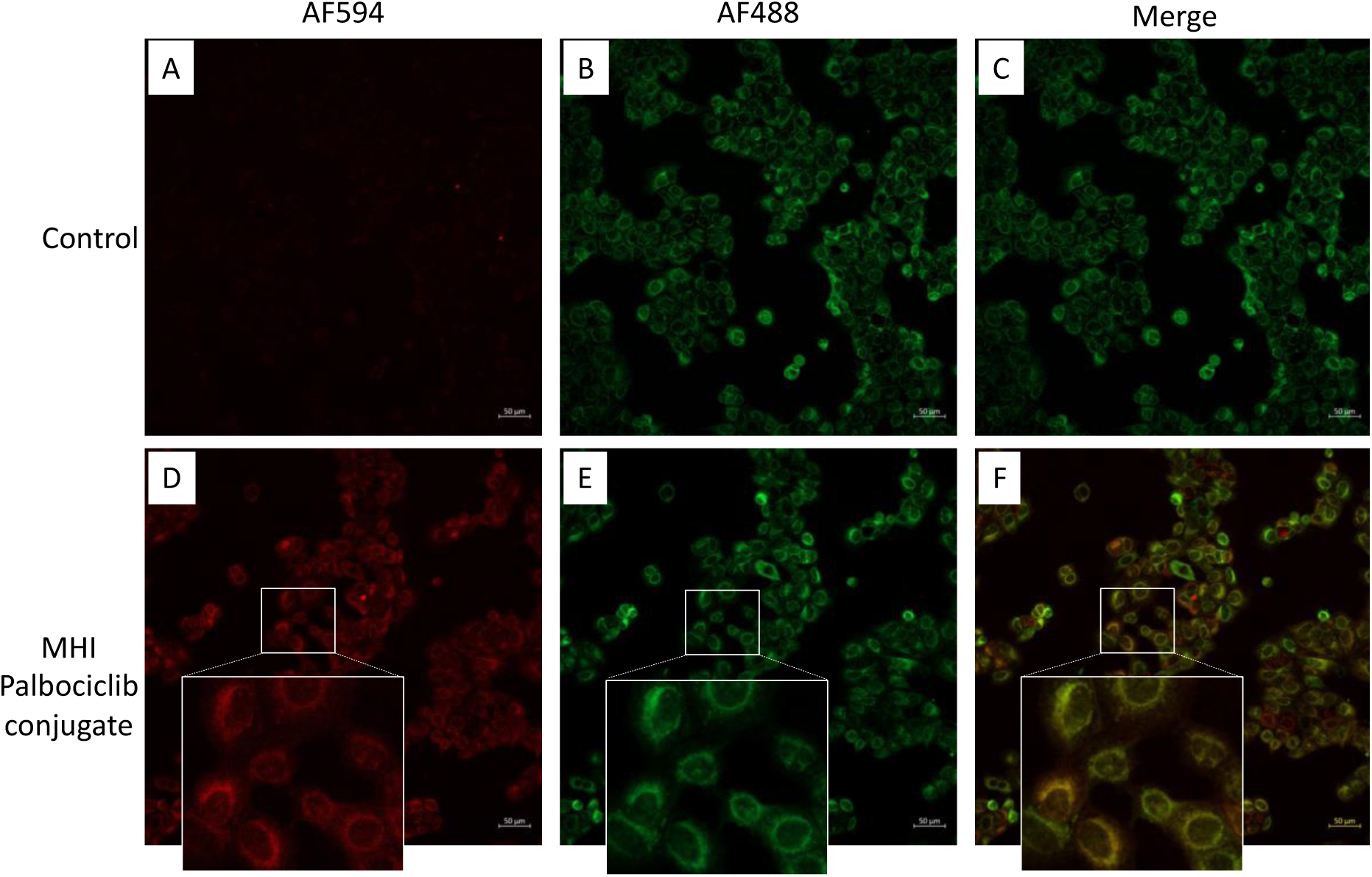
MHI-palbociclib localized in mitochondria. Red fluorescence (MHI-palbociclib) **(A, D**) and green fluorescence (Mitotracker Green) **(B, E)** were captured using confocal microscopy. DMSO was used as control. **(F)** Merged image showing MHI-palbociclib (red) co-localized in mitochondria (Mitotracker Green). Scale bar: 50 μm.

### Chemical Space

The calculated molecular descriptors MW (molecular weight), log *P* (water-octanol partition coefficient), HD (hydrogen bond donors), HA (hydrogen bond acceptors) and solvent accessible surface (SAS) area are given in Table 2 derived using the Scigress program.^25^

**Table 2.** The molecular descriptors as calculated by the software Scigress

|  | MW<br>(g/mol) | Log<br>P | HD | HA | SAS<br>(Å<sup>2</sup>) |
| --- | --- | --- | --- | --- | --- |
| **Palbociclib** | 447.5 | 2.3 | 2 | 9 | 451.9 |
| **MHI148** | 684.3 | 7.7 | 2 | 5 | 641.6 |
| **MHI-palbociclib** | 1543.4 | 11.2 | 2 | 21 | 1198.4 |

Large differences in molecular weights, Log P and SAS area are observed between MHI-palbociclib, MHI-148 and palbociclib as seen in Table 2. MHI-palbociclib has obviously much larger mass, its log P value is >10 characterizing it as extremely lipophilic, and the SAS area is much larger. The lipophilic MHI moieties in which the positive charge is dispersed over a large surface area can easily pass through lipid bilayers and the potential gradient drives their accumulation into the mitochondrial matrix. The calculated properties indicate that the distribution of palbociclib in cells will likely be different to that of MHI-palbociclib since the lipophilic conjugate can be expected to be present in lipophilic environments such as the mitochondria.^38^

## Discussion

We have synthesized MHI-palbociclib according to the previously reported protocol and demonstrated that MHI-palbociclib showed increased potency in inhibiting cell growth and viability as compared to palbociclib in breast cancer cell lines MCF-7 and MDA-MB-231 (and also in non-cancerous HEK293, 51D1 and 51D1.3 cells). In MHI-palbociclib treated cells, the increased cytotoxic effect (as measured by PI staining in Figure 5) presumably contributed to the observed efficacy in growth inhibition and viability reduction (Figure 2F and I), whilst the cells treated with palbociclib showed the characteristic and expected cytostatic effects^3,39^ (minimal PI staining, no effect on viability and growth, and increased cell cycle G1 arrest) in our study. Our results suggests that MHI-palbociclib has a different mechanism of action to that of palbociclib.

Previous studies have demonstrated the preferential retention of NIRF dyes in the mitochondria of cancer cell lines,^12^ while palbociclib was retained in the lysosomal compartment, presumably because of protonation of the piperazine side chain in this acidic organelle.^40^ The localization of MHI-palbociclib in mitochondria (Figure 6) recapitulated that of the dye MHI-148 rather than that of palbociclib, which is not unexpected given that in the MHI-palbociclib, the basic piperazine group is converted to a neutral amide functionality. However, we cannot exclude the possibility that some conjugate may have undergone hydrolysis to release palbociclib. The chemical space data indicated that MHI-palbociclib is very lipophilic (its log P value is >10) and could therefore be expected to accumulate in the mitochondria, supporting the localization in mitochondria observed in our study. While the palbociclib moieties of MHI-palbociclib might retain their kinase inhibitory properties when conjugated, intracellular kinase inhibition could be altered by the lipophilic nature of MHI-palbociclib and by its sequestration in mitochondria.

The significant difference in G1 cell cycle arrest induction in palbociclib-treated cells compared to MHI-palbociclib-treated cells (Figure 4) demonstrated that the MHI-palbociclib indeed had a different mode of action compared to the parent compound palbociclib, at least in the cell lines we examined.

It is known that serum albumin accumulates in regions of proliferating tumor cells, which is thought to be a result of the enhanced permeability and retention effect (EPR).^41^ As a natural ligand carrier, albumin has shown remarkable promise as a carrier for anti-cancer agents to promote their accumulation within tumors.^41^ In the presence of albumin, MHI-148 rapidly forms a non-covalent complex, and, like other HMCDs containing a *meso*-chlorine, over time the albumin free thiol displaces the *meso*-chlorine to form a covalent adduct.^13^ As the cell-based assays in our study were performed in the presence of 5% FBS (equating to an albumin concentration of ∼19 μM in the culture medium), the albumin concentration was in excess compared to the concentration of MHI-148 and MHI-palbociclib in all experiments. Therefore the cell-based assays involving MHI-148 and MHI-palbociclib presumably contained a mixture of species including the parent drug, a noncovalent adduct, and a covalent adduct. It is highly likely that MHI-148 will enhance cancer cell accumulation of derived conjugates. However, the role of albumin-binding in intracellular trafficking remains to be investigated thoroughly and is likely dependent on both the type of therapeutic cargo and formulation.

From our study, MHI-palbociclib is likely to engage a different target and perform a different function other than CDK4/6 inhibition. It is worth noting that in MHI-palbociclib, palbociclib was linked to MHI-148 by an amide bond, which can be expected to be much more stable intracellularly than the ester group used to link dasatinib to MHI-148.^15^

For future study, the use of a cleavable linker such as a disulfide-based linker ^42^ may allow for selective intracellular release of the active molecule (palbociclib) upon glutathione reduction and linker cleavage, since many cancer cells contained elevated GSH levels in comparison with normal cells.^43^ Since MHI-palbociclib was localized to mitochondria, a monosubstituted disulfide linker could be considered as the chemical cleavage of disulfide linkers in the mitochondria was reported to differ significantly.^44^ Another important aspect is to maintain the tumor targeting potential of the conjugate to achieve good therapeutic index for future development.

## Conflict of Interest

The authors have no conflicts of interest to disclose.

## Acknowledgments

The authors would like to acknowledge the support of the Auckland Cancer Society Research Centre at the University of Auckland. EL acknowledges support from the New Zealand Breast Cancer Foundation Belinda Scott Science Fellowship. PT acknowledges support of the School of Medical Sciences and Auckland Cancer Society Research Centre at the University of Auckland. MT acknowledges the support of the Cancer Society Auckland-Northland Division. TP acknowledges support from the Douglas Charitable Trust. JJ, PC, TP, EC acknowledge the support of Cancer Research Trust New Zealand (CRTNZ 2013 RPG), Maurice Phyllis Paykel Trust (203123) and the support of the Neurological Foundation New Zealand. The funders had no role in the study design, data collection, analysis and interpretation, publication decision or writing of the article.

---

**Supplementary Table S1.**
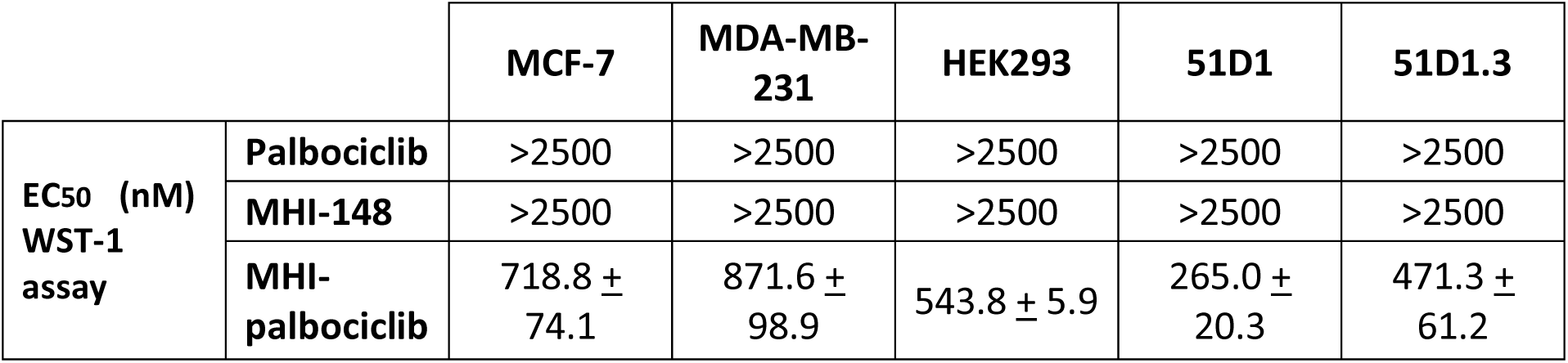
EC_50_ values of palbociclib, MHI-148 and MHI-palbociclib in breast cancer MCF-7 and MDA-MB-231 cell lines, and non-cancerous HEK293, 51D1 and 51D1.3 cell lines using WST-1 assay.

## Notes

### Competing Interest Statement

The authors have declared no competing interest.

## References

1. Dai, X., et al. Breast cancer intrinsic subtype classification, clinical use and future trends. Am J Cancer Res 5, 2929–2943 (2015).

2. Hanahan, D. & Weinberg, R.A. The hallmarks of cancer. Cell 100, 57–70 (2000).

3. Trotter, E.W. & Hagan, I.M. Release from cell cycle arrest with Cdk4/6 inhibitors generates highly synchronized cell cycle progression in human cell culture. Open Biol 10, 200200 (2020).

4. Otto, T. & Sicinski, P. Cell cycle proteins as promising targets in cancer therapy. Nature reviews. Cancer 17, 93–115 (2017).

5. Wedam, S., et al. FDA Approval Summary: Palbociclib for Male Patients with Metastatic Breast Cancer. Clinical cancer research : an official journal of the American Association for Cancer Research 26, 1208–1212 (2020).

6. Roncato, R., et al. CDK4/6 Inhibitors in Breast Cancer Treatment: Potential Interactions with Drug, Gene, and Pathophysiological Conditions. International journal of molecular sciences 21(2020).

7. Hafeez, U., Parakh, S., Gan, H.K. & Scott, A.M. Antibody-Drug Conjugates for Cancer Therapy. Molecules 25(2020).

8. Hapuarachchige, S. & Artemov, D. Theranostic Pretargeting Drug Delivery and Imaging Platforms in Cancer Precision Medicine. Frontiers in oncology 10, 1131 (2020).

9. Shi, C., Wu, J.B. & Pan, D. Review on near-infrared heptamethine cyanine dyes as theranostic agents for tumor imaging, targeting, and photodynamic therapy. J Biomed Opt 21, 50901 (2016).

10. Cooper, E., et al. The Use of Heptamethine Cyanine Dyes as Drug-Conjugate Systems in the Treatment of Primary and Metastatic Brain Tumors. Frontiers in oncology 11, 654921 (2021).

11. Choi, P.J., et al. Heptamethine Cyanine Dye Mediated Drug Delivery: Hype or Hope. Bioconjug Chem 31, 1724–1739 (2020).

12. Yang, X., et al. Near IR Heptamethine Cyanine Dye–Mediated Cancer Imaging. Clinical Cancer Research 16, 2833–2844 (2010).

13. Thavornpradit, S., Usama, S.M., Lin, C.M. & Burgess, K. Protein labelling and albumin binding characteristics of the near-IR Cy7 fluorophore, QuatCy. Org Biomol Chem 17, 7150–7154 (2019).

14. Usama, S.M., Zhao, B. & Burgess, K. Fluorescent kinase inhibitors as probes in cancer. Chem Soc Rev (2021).

15. Usama, S.M., Jiang, Z., Pflug, K., Sitcheran, R. & Burgess, K. Conjugation of Dasatinib with MHI-148 Has a Significant Advantageous Effect in Viability Assays for Glioblastoma Cells. ChemMedChem 14, 1575–1579 (2019).

16. Wu, J.B., et al. Near-infrared fluorescence heptamethine carbocyanine dyes mediate imaging and targeted drug delivery for human brain tumor. Biomaterials 67, 1–10 (2015).

17. Yang, X., et al. Near IR heptamethine cyanine dye-mediated cancer imaging. Clinical cancer research : an official journal of the American Association for Cancer Research 16, 2833–2844 (2010).

18. Xiao, L., et al. Heptamethine cyanine based (64)Cu-PET probe PC-1001 for cancer imaging: synthesis and in vivo evaluation. Nucl Med Biol 40, 351–360 (2013).

19. Hinz, J.M., et al. Repression of mutagenesis by Rad51D-mediated homologous recombination. Nucleic acids research 34, 1358–1368 (2006).

20. Leung, E.Y., et al. Derivation of Breast Cancer Cell Lines Under Physiological (5%) Oxygen Concentrations. Frontiers in oncology 8, 425 (2018).

21. Vichai, V. & Kirtikara, K. Sulforhodamine B colorimetric assay for cytotoxicity screening. Nat Protoc 1, 1112–1116 (2006).

22. Leung, E.Y., et al. Endocrine Therapy of Estrogen Receptor-Positive Breast Cancer Cells: Early Differential Effects on Stem Cell Markers. Frontiers in oncology 7, 184 (2017).

23. Leung, E., et al. Validating TDP1 as an Inhibition Target for the Development of Chemosensitizers for Camptothecin-Based Chemotherapy Drugs. Oncol Ther (2021).

24. Reynisson, J., et al. Evidence that phospholipase C is involved in the antitumour action of NSC768313, a new thieno[2,3-b]pyridine derivative. Cancer Cell Int 16, 18 (2016).

25. Scigress Ultra V. F.J 2.6. (2008–2016).

26. Allinger, N.L., Yuh, Y.H. & Lii, J.H. Molecular mechanics. The MM3 force field for hydrocarbons. 1. Journal of the American Chemical Society 111, 8551–8566 (1989).

27. Lii, J.H. & Allinger, N.L. Molecular mechanics. The MM3 force field for hydrocarbons. 2. Vibrational frequencies and thermodynamics. Journal of the American Chemical Society 111, 8566–8575 (1989).

28. Lii, J.H. & Allinger, N.L. Molecular mechanics. The MM3 force field for hydrocarbons. 3. The van der Waals’ potentials and crystal data for aliphatic and aromatic hydrocarbons. Journal of the American Chemical Society 111, 8576–8582 (1989).

29. Gotō, H. & Ōsawa, E. An efficient algorithm for searching low-energy conformers of cyclic and acyclic molecules. Journal of the Chemical Society, Perkin Transactions 2, 187–198 (1993).

30. Ghose, A.K. & Crippen, G.M. Atomic physicochemical parameters for three-dimensional-structure-directed quantitative structure-activity relationships. 2. Modeling dispersive and hydrophobic interactions. Journal of Chemical Information and Computer Sciences 27, 21–35 (1987).

31. Klamt, A. & Schüürmann, G. COSMO: a new approach to dielectric screening in solvents with explicit expressions for the screening energy and its gradient. Journal of the Chemical Society, Perkin Transactions 2, 799–805 (1993).

32. Chen, P., et al. Spectrum and Degree of CDK Drug Interactions Predicts Clinical Performance. Molecular cancer therapeutics 15, 2273–2281 (2016).

33. Cullen, A., et al. Experimental crystal structure determination of MHI-148. Cambridge Crystallographic Data Centre (2021).

34. Finn, R.S., et al. PD 0332991, a selective cyclin D kinase 4/6 inhibitor, preferentially inhibits proliferation of luminal estrogen receptor-positive human breast cancer cell lines in vitro. Breast cancer research : BCR 11, R77 (2009).

35. Cretella, D., et al. Pre-treatment with the CDK4/6 inhibitor palbociclib improves the efficacy of paclitaxel in TNBC cells. Scientific reports 9, 13014 (2019).

36. Franco, J., Balaji, U., Freinkman, E., Witkiewicz, A.K. & Knudsen, E.S. Metabolic Reprogramming of Pancreatic Cancer Mediated by CDK4/6 Inhibition Elicits Unique Vulnerabilities. Cell Rep 14, 979–990 (2016).

37. Pietkiewicz, S., Schmidt, J.H. & Lavrik, I.N. Quantification of apoptosis and necroptosis at the single cell level by a combination of Imaging Flow Cytometry with classical Annexin V/propidium iodide staining. J Immunol Methods 423, 99–103 (2015).

38. Murphy, M.P. Targeting lipophilic cations to mitochondria. Biochimica et biophysica acta 1777, 1028–1031 (2008).

39. Lypova, N., et al. Targeting Palbociclib-Resistant Estrogen Receptor-Positive Breast Cancer Cells via Oncolytic Virotherapy. Cancers (Basel) 11(2019).

40. Llanos, S., et al. Lysosomal trapping of palbociclib and its functional implications. Oncogene 38, 3886–3902 (2019).

41. Hoogenboezem, E.N. & Duvall, C.L. Harnessing albumin as a carrier for cancer therapies. Advanced drug delivery reviews 130, 73–89 (2018).

42. Li, S., et al. Targeted Methotrexate Prodrug Conjugated With Heptamethine Cyanine Dye Improving Chemotherapy and Monitoring Itself Activating by Dual-Modal Imaging. Frontiers in Materials 5(2018).

43. Perry, R.R., Mazetta, J.A., Levin, M. & Barranco, S.C. Glutathione levels and variability in breast tumors and normal tissue. Cancer 72, 783–787 (1993).

44. Lei, E.K. & Kelley, S.O. Delivery and Release of Small-Molecule Probes in Mitochondria Using Traceless Linkers. J Am Chem Soc 139, 9455–9458 (2017).

